# Behavioral state-dependent noradrenergic dynamics in the primary somatosensory and prefrontal cortices during tactile detection tasks

**DOI:** 10.64898/2026.01.16.699887

**Authors:** Craig Kelley, Tim Lantin, Cody Slater, Marc Sorrentino, Kunpeng Yu, Bo Yuan, Michael Kann, Yuxiang Liu, Jung Soo Kim, Qi Wang

**Author notes:** Contact Info: Correspondence should be addressed to: Professor Qi Wang, Department of Biomedical Engineering Columbia University, ET 351, 500 W. 120^th^ Street, New York, NY 10027.

## Abstract

Animals must integrate sensory information, ignore behaviorally irrelevant stimuli, and respond to behaviorally relevant stimuli to find food, find mates, avoid predators, and ultimately survive. In mammals, these goal-directed behaviors require the coordinated activity of many brain regions, including sensory and prefrontal cortices and neuromodulatory brainstem nuclei like the locus coeruleus (LC), which is the brain’s primary source of norepinephrine (NE). NE release resulting from LC activity and arousal indexed by pupil size exert strong influences on goal-directed behavior. We explored the relationships between pupil size, cortical noradrenergic dynamics, and behavior in a tactile signal detection task. We monitored pupil dynamics and fluorescent GRAB_NE_ signals in somatosensory and medial prefrontal cortices simultaneously during task execution and found that pupil size and synchronization of GRAB_NE_ signals at baseline were strong predictors of whether animals chose to respond. Baseline and post-reward cortical GRAB_NE_ levels varied strongly with pupil-linked arousal. We also employed a generalized linear model - hidden Markov model (GLM-HMM) framework to identify distinct, stable behavioral states throughout the task that characterize task performance. We found distinct psychometric curves, task-related pupil dynamics, and cortical NE dynamics across these behavioral states.

**Significance Statement:** Behavioral state strongly shapes goal-directed behavior, which in turn depends on the coordinated activity of distributed brain regions, including the sensory and prefrontal cortices. By simultaneously measuring pupil size and cortical noradrenergic dynamics, and by identifying psychophysically distinct behavioral states during a tactile detection task, this study establishes links between pupil-linked arousal, norepinephrine signaling in somatosensory and prefrontal cortices, and trial outcomes. These findings provide new insight into how the locus coeruleus – norepinephrine system regulates perception and decision-making.

## Introduction

Integrating sensory information, responding to behaviorally relevant stimuli, and ignoring behaviorally irrelevant stimuli is essential to animal survival (Wimmer et al., 2015; Speed and Haider, 2021). Goal-directed behaviors require the coordinated activity of multiple brain regions: thalamus and sensory cortices process and encode stimuli; prefrontal cortex is involved in decision making, planning, and working memory; and neuromodulatory brainstem nuclei dynamically modulate processing in other brain regions through attention and arousal (Pezzulo et al., 2014; Dahl et al., 2022; Cerpa et al., 2023; Jordan and Keller, 2023; Ghosh and Maunsell, 2024; Hansen et al., 2024; Osorio-Forero et al., 2024; Jensen et al., 2025; Kelley et al., 2025). In particular, activity in the locus coeruleus, the brain’s primary source of norepinephrine (NE), has been shown to modulate feedforward processing of behaviorally-relevant stimuli and correlate with performance in goal-directed behaviors (Devilbiss et al., 2006; de Gee et al., 2017; Clewett et al., 2018; Totah et al., 2018; Vazey et al., 2018; Rodenkirch et al., 2019; Orlando et al., 2023; Grimm et al., 2024; Rodenkirch and Wang, 2024). Recent developments in genetically encoded fluorescent sensors *in vivo* have opened a window into the dynamics of NE released in brain regions targeted by LC projections during sensory and cognitive processing, learned behaviors, and sleep (Feng et al., 2019; Breton-Provencher et al., 2022; Kjaerby et al., 2022; Osorio-Forero et al., 2024; Liu et al., 2025).

Pupil diameter is a reliable physiological biomarker of arousal and predictor of task performance in perceptual and cognitive tasks (Hess and Polt, 1960; Kahneman and Beatty, 1966; Nassar et al., 2012; de Gee et al., 2014; Reimer et al., 2014; Ebitz and Platt, 2015; McGinley et al., 2015; Vinck et al., 2015; Urai et al., 2017; Schriver et al., 2020; Narasimhan et al., 2023). There is also strong evidence of a link between neural activity in LC and dynamic changes in pupil size; however, the relationships between LC activity, pupil size, and arousal are strongly nonlinear and nonexclusive (Joshi et al., 2016; Liu et al., 2017). Although large pupil dilations often coincided with bursts of LC firing, reflecting arousal or salient stimuli, whether pupil size alone reliably indexes spiking activity in LC remains hotly debated (Megemont et al., 2022).

During goal directed behaviors, performance can vary across distinct behavioral states underpinned by distinct neural dynamics (Akella et al., 2025). Ashwood et al. (2022) developed a method using hidden Markov models (HMMs) to represent discrete behavioral states, and each state was associated with a generalized linear model (GLM) characterizing psychometric performance. Previous behavior, measures of pupil-linked arousal, uninstructed movements, and neural activity can all predict changes in behavioral state (Ashwood et al., 2022; Hulsey et al., 2024; Akella et al., 2025; Johnson et al., 2025; Yin et al., 2025).

Here we explored the relationships between pupil size fluctuations, noradrenergic dynamics, and behavioral state in a tactile stimulus detection task. Mice were trained to respond to behaviorally relevant tactile signals in pursuit of water rewards. We simultaneously monitored pupil dynamics and GRAB_NE_ signals in somatosensory and medial prefrontal cortices during the task. GRAB_NE_ is a genetically encoded fluorescent sensor that responds to the local release of NE (Feng et al., 2019), and we used fiber photometry to measure regional release of NE during behavior. We found that baseline pupil size and baseline synchronization of GRAB_NE_ signals, but not baseline cortical GRAB_NE_ levels, were strong predictors of whether animals decided to respond. High baseline pupil area was associated with disengagement from the task and reduced cortical GRAB_NE_ signals following reward collection on hit trials. We also employed a generalized linear model - hidden Markov model (GLM-HMM) framework to identify behavioral states throughout the task with state transitions linked to baseline pupil measurements as in Hulsey et al. (2024). The behavioral states identified by the GLM-HMM were psychometrically distinct and temporally stable, and we found distinct noradrenergic dynamics across these behavioral states.

## Methods

### Surgery

All experiments were approved by the Institutional Animal Care and Use Committee (IACUC) at Columbia University and were conducted in compliance with NIH guidelines. Mice (B6, Jax #: 000664) underwent survival surgery for AAV injection and implantation of optical fibers and head plate at age of 3-6 months. Mice were initially anesthetized with isoflurane in oxygen (5% induction, 2% maintenance) and then secured in a stereotaxic frame. Body temperature was maintained at ∼37 ℃ using a feedback-controlled heating pad (FHC, Bowdoinham, ME). After the mouse’s condition stabilized but before an incision was made on the scalp, lidocaine hydrochloride and buprenorphine (0.05 mg/kg) were administered subcutaneously to ensure analgesics were on board during the whole surgery. To measure cortical NE dynamics during tactile signal detection tasks, AAVs encoding GRAB_NE_ (AAV9-hSyn-NE2h, WZ Biosciences) were injected into the mPFC (AP: +2.3 mm, ML: 1.2 mm, DV: -2.0 mm) and S1 (AP: -1.30 mm, ML: +3.3 mm, DV: -0.72). Before the injection, small burr holes were drilled above the mPFC and S1 and saline was applied to each craniotomy to prevent exposed brain surface from drying out. Pulled capillary glass micropipettes were back-filled with AAV solution, which was subsequently injected into the target brain regions (∼200 nL each site) at 0.8 nL/s using a precision injection system (Nanoliter 2020, World Precision Instruments, Sarasota, FL). The micropipette was left in place for at least 10 minutes following each injection and then slowly withdrawn. Following GRAB_NE_ AAV injection, an optical fiber (200 μm diameter, NA = 0.39) was implanted with the tip of the fiber placed approximately 0.15 mm above the injection site. Metabond (Parkell Inc., Edgewood, NJ) was used to build a headcap to bond the fibers. The ferrules and headplate were then cemented in place with dental acrylic. At the conclusion of the surgery, Baytril (5 mg/kg) and Ketoprofen (5 mg/kg) were administered. Four additional doses of Baytril and two additional doses of Ketoprofen were provided every 24 hours after the surgery day. Animals’ weight was measured at least once per day for 5 days.

### Fiber photometry recording

Fiber photometry recording was performed approximately 3 weeks following surgery to allow sufficient time for viral expression. Fluorescence signals mediated by the GRAB_NE_ sensors were recorded using a 2-channel fiber photometry system (Doric Lenses). For each channel, the excitation light with 465 nm wavelength was generated by an LED (CLED_465, Doric Lenses) and passed through a MiniCube (iFMC4_AE (405)E(460–490)_F(500–550)_S, Doric Lenses). Emission fluorescence from the GRAB_NE_ sensor was measured by an integrated PMT detector in the MiniCube. The fiber photometry recordings were run in a ‘Lock-in’ mode controlled by Doric Neuroscience Studio (V5.4.1.12), where the intensity of the excitation lights was modulated at frequencies of 208.62 Hz and 572.21 Hz to avoid contamination from other light sources in the room and crosstalk between the excitation lights (Liu et al., 2025). The demodulated signal processed by the Doric fiber photometry console was low-pass filtered at 25 Hz and sampled at 12 kHz with a 16-bit ADC. The fiber photometry system was synchronized with the behavioral apparatus through TTLs generated by the xPC target real-time system (MathWorks, Massachusetts). All photometry data were decimated to 120 Hz by Doric Neuroscience Studio software and saved for offline analysis. We used signal background estimation with a 3 s moving window to remove drifts in the baseline signal while preserving peaks and oscillations (MATLAB function *msbackadj*).

### Pupillometry

We monitored mouse pupils continuously during the task. Pupil recordings were obtained using a custom pupillometry system (Schriver et al., 2018). The camera captured images at 10 frames per second via TTL triggers from the xPC real-time system. Pupil images were streamed to a high-speed solid-state drive for offline analysis. As previously described, the DeepLabCut toolbox was used for automated segmentation of the pupil contour (Weiss et al., 2025). Elliptical regression was then applied to fit the labeled points, enabling the computation of pupil size based on the fitted contour. To ensure segmentation accuracy, approximately 5% of segmented images were randomly selected and inspected. Pupil size during periods of blinks was estimated by interpolating over frames immediately preceding and following the blinks. If DeepLabCut did not recognize pupil contour due to either poor video quality or animal’s eyelid covering a significant portion of pupil in >33% of the recorded video frames, the session was excluded from pupillometry analysis. Prior to further analysis, a fourth-order non-causal low-pass filter with a cutoff frequency of 3.5 Hz was applied to the pupil size data (Joshi et al., 2016) .

### Behavioral task

Head-fixed mice (n=4; all male; aged 4-7 months) were trained to respond to tactile stimuli applied to their whiskers via air puffs by licking a water spout. Mice were singly housed on a 12-hour light-dark cycle with ad libitum access to food. Mice were water restricted to maintain ≥ 85% of their body weight at 3 weeks following surgery. Prior to training, mice were acclimated to head-fixation for 10 minutes a day for 2 days. Mice were trained in stages: first with two sessions of classical conditioning (water reward paired with tactile stimuli regardless of response), two sessions of operant conditioning (mice must lick the reward spout in response to tactile stimuli to receive water reward; no distractor puffs), followed by the full task (see below). Mice needed to complete 3 consecutive sessions with d’ > 1 before sessions were included in analysis. Mice remained head-fixed for the duration of the session. During a session, each trial began with an onset tone followed by a delay period lasting 4-8 s. During the delay period, 1-3 distractor air puffs (not aimed at the mouse or its whiskers, but close enough to hear) were delivered at random times. Responses to a distractor stimulus (licking the water spout within 500 ms of the distractor) resulted in restarting the trial. The mouse needed to withhold licks for the last 2 s of the delay period in order for a stimulus to be delivered. On 40% of trials, no stimulus (0 PSI) was delivered (i.e. catch trials). On the remaining 60% of trials, air puffs were delivered to left whiskers at different pressures (0.5 – 20 PSI) with equal probability. Stimulus delivery, which lasted 200 ms, was followed by a 500 ms window of opportunity, during which the mouse could respond to the tactile stimulus by licking the water spout. The distance between the nozzle delivering the air puff and the platform where the mouse performed the task was fixed across session, as was the orientation of the nozzle. Responses to tactile stimuli within the window of opportunity were rewarded with 2.3 µL of sweetened water (10% sucrose) followed by a 2 s cool down period before the start of the next trial. Responses to 0 PSI stimuli on catch trials resulted in a 15 s timeout before the start of the next trial. Sessions lasted 129 ± 5 trials (56.4 ± 0.1 mins). We recorded between 6 and 17 sessions from each animal for 49 sessions in total.

### Histology

Since GRAB_NE_ constructs use a GFP-based fluorescent reporter, we confirmed expression of GRAB_NE_ at the end of the study using immunofluorescence. Mice were transcardially perfused with PBS followed immediately by ice-cold 4% paraformaldehyde. The brain was removed carefully and post-fixed overnight at 4 °C in 4% paraformaldehyde, and then cryopreserved in a 30% sucrose (wt/vol) in PBS solution for 3 days at 4 ℃. Brains were embedded in Optimum Cutting Temperature Compound, and 30-μm coronal slices were sectioned using a cryostat. Brain slices were washed 4x in PBS and then incubated in 10% normal goat serum contained with 0.5% Triton X-100 in PBS for 2 hours. This was followed by primary antibody incubation overnight at room temperature using a chicken anti-GFP primary antibody (1:500). On the next day, slices were washed 3x in PBS + Tween (0.0005%) solution followed by secondary antibody incubation for 2 hours at room temperature using an Alexa Fluor 488-conjugated goat anti-chicken (1:800). The slices were then washed 3x in PBS + Tween solution and 1x with PBS only followed by coverslipping using Fluoromount-G medium with DAPI. Slices were imaged using 8X objective in a slide scanner (Nikon AZ100) for verification of AAV transfection.

### Analyses

All analyses were performed using custom code written in MATLAB (Mathworks, Natick, MA) and Python available at https://github.com/Neural-Control-Engineering/Neurodynamic-control-toolbox. All pupillometry and fiber photometry recordings from each session were z-scored prior to further analyses. Baseline measurements from pupillometry and fiber photometry recordings were averaged over 0.5 s prior to stimulus delivery. The magnitude of stimulus-induced pupil dilations and increases in NE signal were computed by subtracting the baseline value from the maximum pupil size or from GRAB_NE_ signal over the 6 s following stimulus delivery on each trial. When computing cross correlations between GRAB_NE_ in S1 and mPFC, we used 4 s of baseline activity.

We employed the generalized linear model - hidden Markov model (GLM-HMM) framework developed by Ashwood et al. (2022) with the pupil-driven state transitions introduced by Hulsey et al. (2024) to jointly predict responses to tactile stimuli and identify latent behavioral states. We first fit a global model on data pooled across animals, then used its parameters to initialize individual per-mouse fits with K = 2, 3, and 4 states. Stimulus strength and a bias term served as observation-model predictors of the go/no-go decision (response vs. withheld), and baseline pupil area drove the transitions between states. Cross-validation was performed at the session level: sessions were randomly partitioned into five folds (shuffled with fixed random seed), and models were fit on four folds and evaluated on the held-out fold. We selected K = 3 based on held-out log-likelihood and report all downstream analyses on the three-state models. As a baseline, we compared GLM-HMM performance to a classical lapse model (Prins, 2012; Ashwood et al., 2022).

### Statistics

All plotted metrics were performed by computing session averaged metrics (e.g. average baseline pupil area on hit trials for each session) and plotting the mean and standard error of those sessions averaged metrics unless noted otherwise. Statistical comparisons were also performed at the level of these session-averaged metrics unless otherwise noted. Statistical tests between pairs of conditions were two-sided. A one-sample Kolmogorov-Smirnov test was first used to assess the normality of data before performing statistical tests. If the samples were normally distributed, a paired or unpaired t-test was used. Otherwise, the two-sided Mann-Whitney U-test was used for unpaired samples or the two-sided Wilcoxon signed-rank test for paired samples. Bonferroni correction was used for multiple comparisons. Analysis of variance (ANOVA) was used to evaluate differences across multiple factors/conditions (e.g. trial outcome). Repeated measures ANOVA was used to account for measurements from the same session or animal.

## Results

### Behavioral performance in tactile detection tasks

To understand behaviorally relevant NE dynamics in the medial prefrontal cortex (mPFC) and the barrel cortex (S1), we trained 4 mice to perform tactile detection tasks (Stuttgen and Schwarz, 2008; Ollerenshaw et al., 2012). In the task, the animals were required to lick within the window of opportunity (500 ms) to indicate the perception of whisker stimulation induced by a brief air puff with different pressures (**Figure 1A, B,** see **Methods**). To gauge false alarm rate, approximately 40% of all trials were catch trials, in which no air puff was delivered. Catch trials where the mouse did not lick during the window of opportunity were classified as correct rejections. Following the presentation of the tactile stimulus, the animals usually licked within approximately 400 ms to indicate the detection of the tactile stimulus. Trials in which the mouse responded within the response window to a genuine tactile stimulation were classified as hits, whereas trials in which the mouse failed to respond were classified as misses. We found that the intensity of tactile stimuli had no effect on reaction time (i.e. time to first lick; **Figure 1C**, ANOVA(stimulus intensity), F(5)=0.75, p=0.58). As expected, response probability increased as the intensity of tactile stimulus increased (**Figure 1D,E**), resulting in a monotonically increasing perceptual sensitivity with stimulus intensity (**Figure 1F**). On hit trials, there was no significant relationship between lick frequency across the trial (5 s following stimulus) and stimulus intensity (ANOVA(stimulus intensity x time), F(5,49)=1.08, p=0.33) with peak lick frequencies following reward delivery and falling off afterward (**Figure 1D)**.

**Figure 1.**
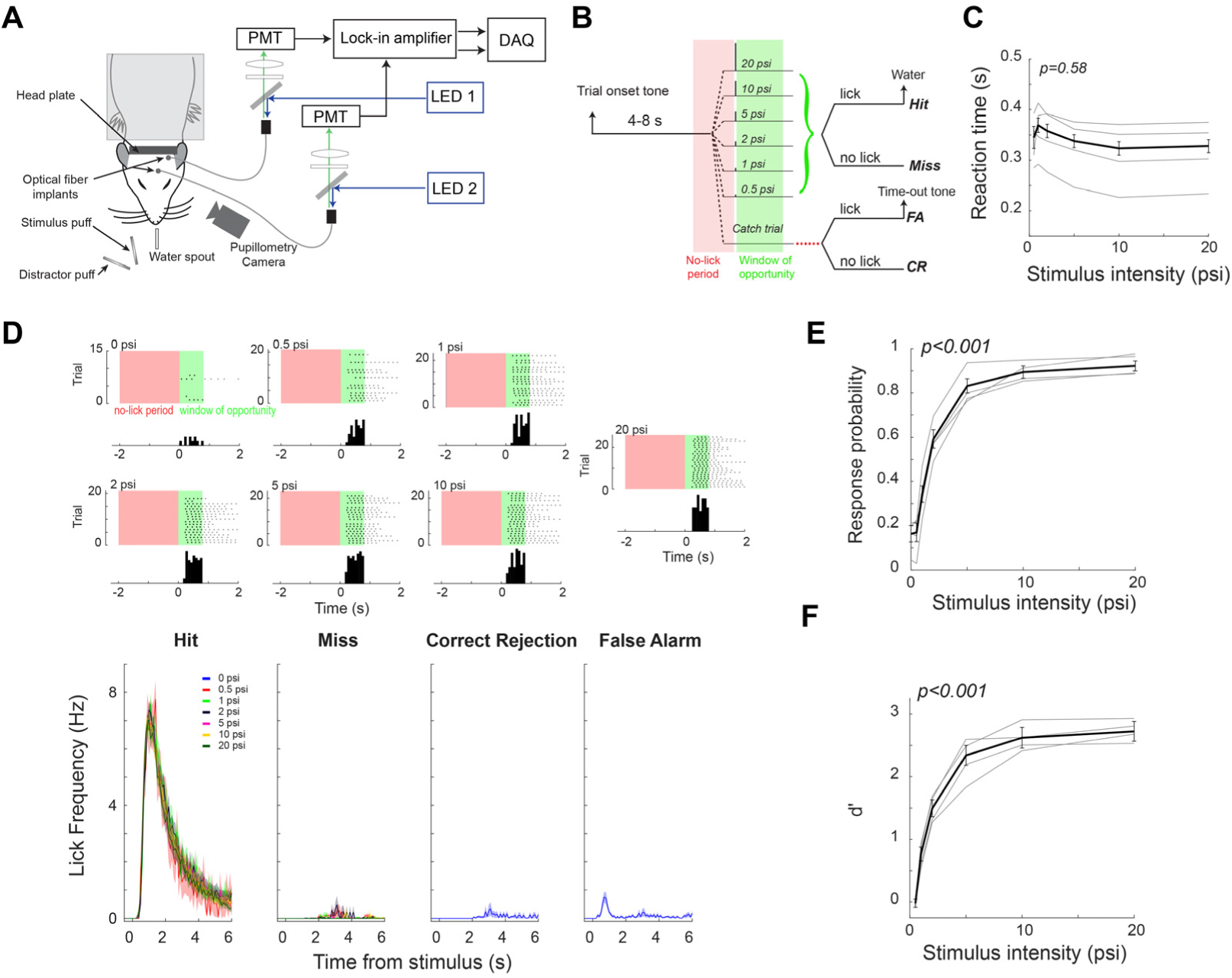
Experimental setup and detection behavior. **A)** Experimental setup. **B)** Diagram of a single trial in the tactile detection task. 0-3 distractor puffs were delivered during the 4-8 s period between trial onset tone and stimulus onset. **C)** Response time for each stimulus intensity. **D)** Top: raster plots from an example session showing responses (licks) to stimuli and each stimulus intensity with histogram of licking response within the window of opportunity below (100 ms non-overlapping bins). Bottom: Average lick frequency across all sessions divided by outcome and stimulus intensity. **E)** Response probability for each tactile stimulus. **F)** Perceptual sensitivity associated with tactile stimuli with different intensities. For C, E, and F, gray lines indicate individual animal averages across sessions, dark line indicates the average across all sessions, and error bars indicate standard error.

### Task-evoked pupil dynamics in whisker detection tasks

We have previously shown that task-evoked phasic pupil dilation reflects different cognitive components in a perceptual decision making task in rats (Schriver et al., 2020), and we hypothesized that pupil dynamics would correlate with behavior similarly in mice. In this study, we imaged pupil size of the mice throughout the behavioral task (z-scored for each session). Similar to rats (Schriver et al., 2018), the pupil of the mice fluctuated throughout the task (**Figure 2A**). The animals’ pupils dilated following the presentation of a tactile stimulus, and the amplitude of task-evoked pupil dilation was positively correlated with the intensity of tactile stimulus when not accounting for trial outcome (**Figure 2B**, ANOVA(stimulus intensity x time), F(6,61)=25.2, p<0.0001). Pupil dynamics around stimulus presentation varied across the four trial outcomes (i.e. hit, correct rejection, miss, and false alarm) with large dilations on response trials (hit and false alarm) and comparatively small dilations, if any, on withheld trials (miss and correct rejection, **Figure 2C**). On hit trials, the dependance of pupil dynamics on stimulus amplitude was weak (ANOVA(stimulus intensity x time), F(5,67)=1.30, p=0.075) while dynamics were strongly influenced by stimulus intensity on miss trials (ANOVA(stimulus intensity x time), F(5,67)=14.4, p<0.0001). Similar to rats’ pupil dynamics in a tactile discrimination task, pupil dilations are dependent upon the baseline pupil size: the larger the baseline pupil size, the smaller the pupil dilation, resulting in negative correlation coefficients between the baseline pupil size and pupil dilation (**Figure 2D-F**). This negative correlation was more profound in response trials than in withheld trials (**Figure 2F**, p<0.001, Wilcoxon signed-rank test).

**Figure 2.**
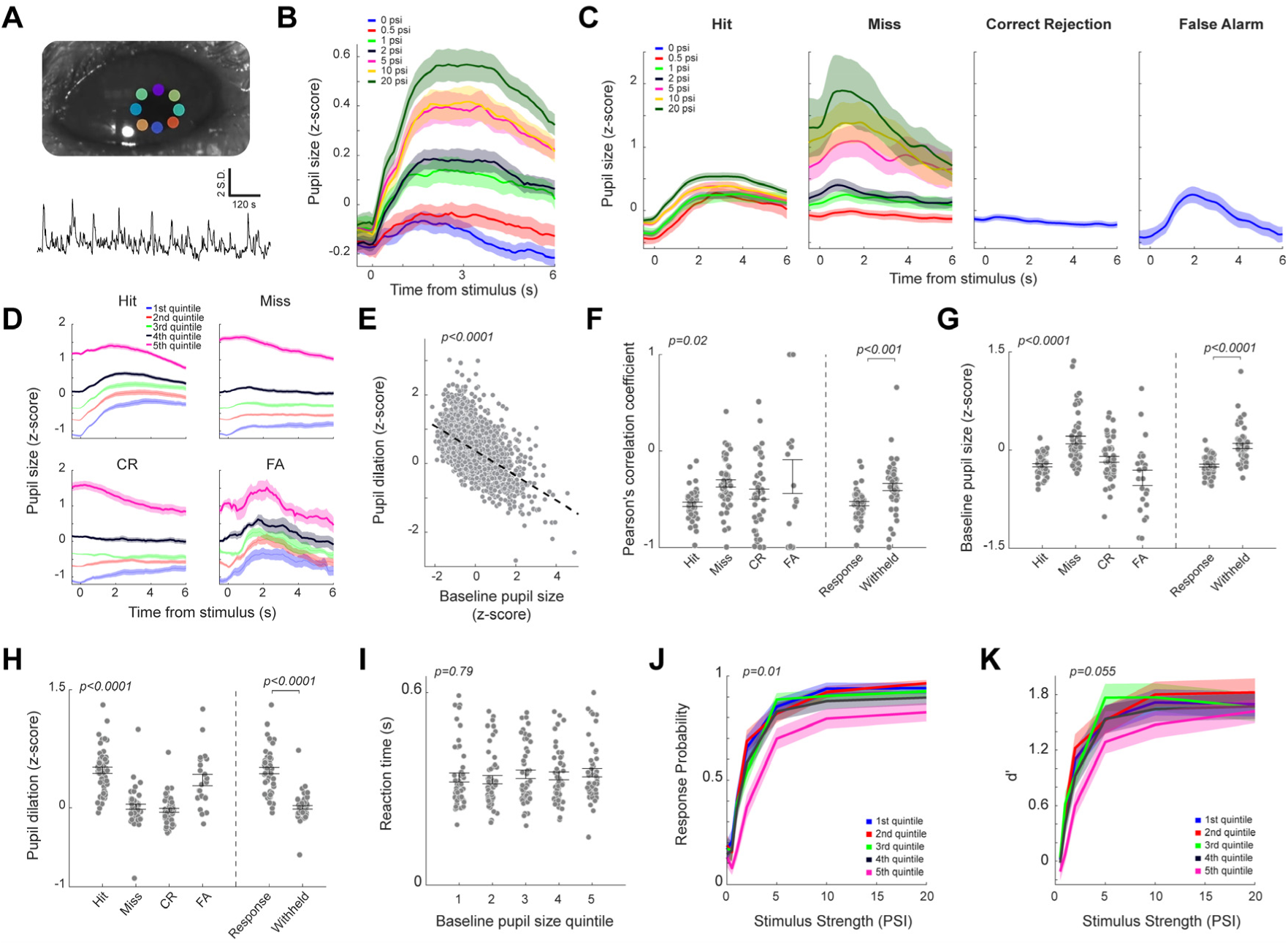
Pupil size fluctuations during the tactile detection task. **A)** Top: Example pupil contour segmented using DeepLabCut. Bottom: Example pupil area fluctuations across a single session. **B)** Pupil responses to tactile stimuli of different intensities regardless of trial outcome. **C)** Pupil dilations associated with the four trial outcomes for each stimulus intensity. **D)** Pupil dilations associated with the four trial outcomes under different baseline pupil size quintiles. **E)** Scatter plot showing the relationship between pupil baseline and pupil dilation across all trials. **F)** Pearson’s correlation coefficients between pupil baseline and dilation for the four trial outcomes (left), and for response and withheld trials (right). **G)** Pupil baseline size for the four trial outcomes (left), and for response and withheld trials (right). **H)** Pupil dilation for the four trial outcomes (left), and for response and withheld trials (right). Baseline-corrected pupil dilations varied significantly across trial outcomes (Extended Data Figure 2-1). **I)** Reaction times for trials in different baseline pupil size quintiles. **J)** Response probability to tactile stimuli of different intensities across trials in different baseline pupil size quintiles. **K)** Perceptual sensitivity to tactile stimuli of different intensities across trials in different baseline pupil size quintiles. Individual data points represent session average metrics unless noted otherwise. Error bars represent standard error of mean.

We found that the baseline pupil size was smallest for false alarm trials (-0.43±0.12), followed by hit (-0.24±0.02), correct rejection (-0.14±0.04), and miss trials (0.16±0.06) (**Figure 2G**, ANOVA(outcome), F(3)=8.65, p<0.0001), suggesting that pupil-linked arousal modulates tactile detection performance. Interestingly, in contrast to rats, the baseline pupil size of the mice was smaller on response trials than on withheld trials (**Figure 2G**, -0.24±0.02 vs. 0.06±0.04, p<0.0001, Wilcoxon signed-rank test, see **Discussion**). However, the task evoked pupil dilation across the four trial outcomes is similar to that in rats performing tactile discrimination tasks, with largest pupil dilation in hit trials (0.48±0.04), followed by false alarm (-0.35±0.07), miss (0.01±0.03), and correct rejection trials (-0.03±0.02) (**Figure 2H**, ANOVA(outcome), F(3)=16.5, p<0.0001). Moreover, task evoked pupil dilation was larger on response trials than on withheld trials (**Figure 2H**, 0.48±0.04 vs. 0.01±0.02, p<0.0001, Wilcoxon signed-rank test). Pupil dilations on hit trials were similar in magnitude to predicted pupil dilations from baseline pupil area, with smaller dilations than predicted on other trial outcomes (**Extended Data Figure 2-1**, ANOVA(outcome), F(3)=32.7, p<0.0001).

To examine the relationship between pupil-linked arousal and behavior, we separated trials by baseline pupil area quintile (1^st^: -2.1 ≤ area < -0.85; 2^nd^: -0.85 ≤ area < -0.50; 3^rd^: -0.50 ≤ area < -0.15; 4^th^: -0.15 ≤ area < 0.47; 5^th^: 0.47 ≤ area ≤ 8.46). Although reaction time did not depend on the baseline pupil size (**Figure 2I**, ANOVA(baseline pupil area quintile), F(4)=0.43, p=0.79), detection probability did vary significantly across pupil size (**Figure 2J**, ANOVA(stimulus intensity x baseline pupil area quintile), F(6,4)=2.80, p=0.01). Detection probability was comparable across most of range of baseline pupil sizes, but there were relative deficits in detection probability when baseline pupil area was highest. There was a similar trend toward a dependence of perceptual sensitivity on baseline pupil size (**Figure 2K**, ANOVA(stimulus intensity x baseline pupil area quintile), F(5,4)=2.17, p=0.055). These data reflected the known left shifted, inverted-U relationship between pupil-linked arousal and task performance with deficits observed on trials with high baseline pupil area, but not low baseline pupil area (McGinley et al., 2015; Beerendonk et al., 2024; de Gee et al., 2024). Even at a much finer binning of trials based on baseline pupil area (e.g. 50 bins), we did not observe significant differences in behavior compared to intermediate baseline pupil area trials (data not shown), which suggests that mice in this study did not enter into states of such high arousal as to result in behavioral deficits.

### Correlated NE dynamics between the primary somatosensory cortex and medial prefrontal cortex

Given the strong association between pupil dynamics and LC dynamics, we next examined NE dynamics in mPFC and S1 during the tactile detection task. Extracellular NE dynamics were measured through fiber photometry recording of fluorescent signals of GRAB_NE_ sensors, which were expressed in the whisker S1 and mPFC (Feng et al., 2019) (**Figure 3A**). GRAB_NE_ signals were z-score for each session. NE signals in the S1 and mPFC prior to stimulus presentation were weakly correlated (**Figure 3B**). We next examined if the correlation between NE dynamics in the S1 and mPFC differed across the four trial outcomes (**Figure 3C**). There was a significant difference in peak correlation coefficient across the trial outcomes (**Figure 3D**, ANOVA(outcome), F(3)=3.13, p=0.03). Correlation between GRAB_NE_ signal in S1 and mPFC were significantly stronger on withheld trials than on response trials (p=0.004, Wilcoxon signed-rank test) but there was no difference in correlation between correct and incorrect trials (**Figure 3D**, p=0.17, Wilcoxon signed-rank test).

**Figure 3.**
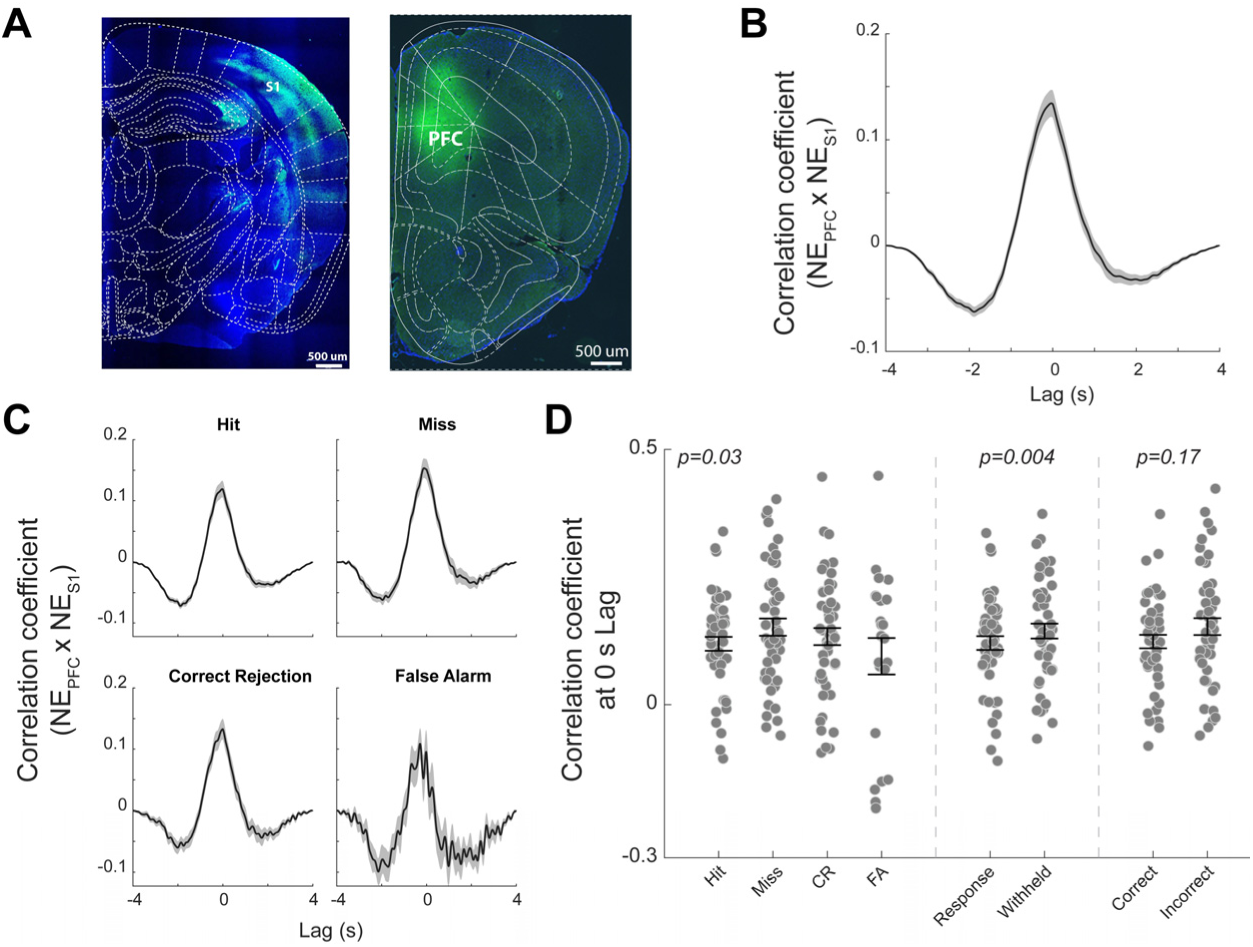
NE dynamics in S1 and mPFC during the tactile detection task. **A)** Immunofluorescence photomicrograph confirming successful expression of GRAB_NE in the barrel cortex (left) and medial prefrontal cortex (right). **B)** Cross-correlation between NE dynamics in S1 and mPFC. **C)** Cross-correlation between pre-stimulus NE dynamics in S1 and mPFC in hit, miss, correct rejection, and false alarm trials. **D)** Session average correlation coefficient at 0 s lag for different trial outcomes. Error bars represent standard error of mean.

### NE dynamics in S1 and mPFC during the detection task

We hypothesized that NE dynamics would differ across the different trial outcomes and stimulus intensities. Like pupil dynamics, task-evoked NE dynamics in S1 and mPFC were dependent on the intensity of tactile stimulus when not accounting for trial outcome (**Figure 4A, G**); however, increases in NE were all-or-none on hit trials, so the dependance on stimulus intensity was a result of psychometric performance, not stimulus amplitude (**Figure 4B, H**). NE dynamics in the S1 were bimodal, with the first peak emerging at around 0.6 s following stimulus presentation. The second peak, which is comparable with the first peak in amplitude, occurred at around 3.5 s following stimulus presentation, well after peak licking activity following reward delivery (**Figure 1D**). Although there were no significant differences in baseline NE levels in S1 across hit, correct rejection, false alarm, and miss trials (ANOVA(outcome), F(3)=1.72, p=0.37), the task evoked increase in S1 NE levels were different across the four trial outcomes (ANOVA(outcome), F(3)=10.0, p<0.0001) (**Figure 4B-D**). In general, NE increases on response trials were higher than on withheld trials (p<0.0001, Wilcoxon signed-rank test) (**Figure 4D**). We next examined if baseline NE levels in S1 had effects on the detection behavior by assessing reaction time and response probability on trials in the different baseline NE quintiles. We found that reaction time was not dependent on baseline NE levels (**Figure 4E**, ANOVA(baseline NE quintile), F(4)=1.11, p=0.35). When separating trials by baseline NE levels in S1 in this manner, we also did not observe any differences in response probability (ANOVA(baseline NE quintile x stimulus intensity), F(4,6)=0.82, p=0.55; **Figure 4F**).

**Figure 4.**
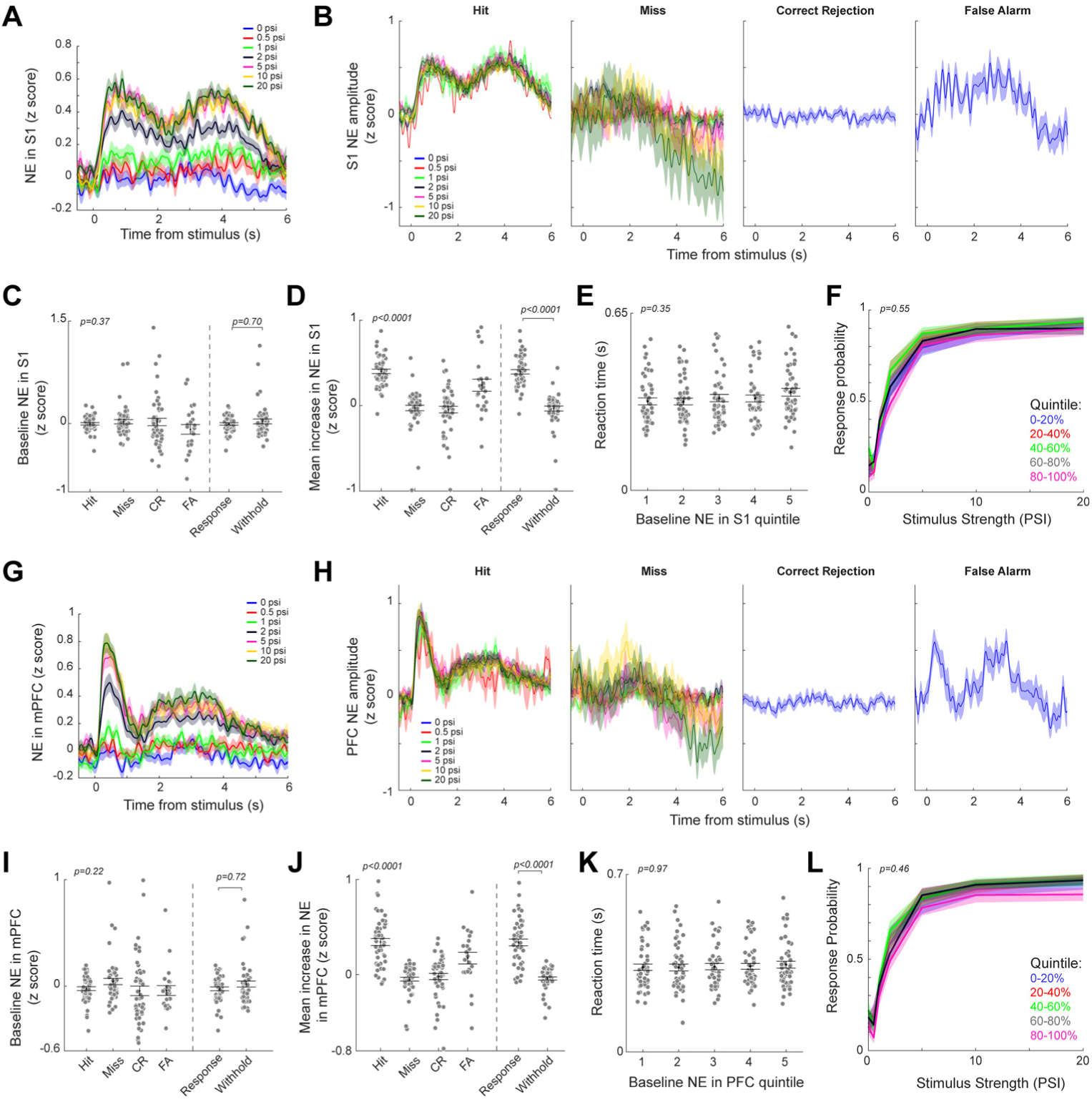
NE dynamics in S1 and mPFC during the tactile detection task. **A)** NE dynamics in S1 evoked by the presentation of different tactile stimuli. **B)** Task evoked NE dynamics in S1 in hit, miss, correct rejection and false alarm trials separated by stimulus intensity. **C)** Baseline NE level in S1 for the four trial outcomes (left), and for response and withheld trials (right). **D)** Mean increase in NE level in S1 for the four trial outcomes (left), and for response and withheld trials (right). **E)** Reaction times during trials in different baseline NE quintiles in S1. **F)** Psychometric curves across trials in different baseline NE quintiles in S1. **G)** NE dynamics in mPFC evoked by the presentation of different tactile stimuli. **H)** Task evoked NE dynamics in mPFC in hit, miss, correct rejection and false alarm trials. **I)** Baseline NE level in mPFC for the four trial outcomes (left), and for response and withheld trials (right). **J)** Mean increase in NE level in mPFC for the four trial outcomes (left), and for response and withheld trials (right). **K)** Reaction times during trials in different baseline NE quintiles in mPFC. **L)** Psychometric curves across trials in different baseline NE quintiles in mPFC. Individual data points represent session average metrics, and error bars represent standard error of mean.

Task-evoked NE dynamics in mPFC were significantly different than those in S1 (ANOVA(region x time), F(1,599)=7.57, p<0.001 across 5 s following stimulus onset on hit trials), but there was no difference in baseline NE levels (p=0.46, Wilcoxon signed-rank test). In mPFC, the decay of the first peak was faster and the amplitude of the second peak was about the half of the first peak (**Figure 4G**), suggesting that the noradrenergic projection to the S1 and mPFC may originate from different subpopulations of LC neurons (see Discussion). Similar to NE dynamics in S1, there were no significant differences in baseline NE levels in mPFC across hit, correct rejection, false alarm, and miss trials (ANOVA(outcome), F(3)=1.46, p=0.22; **Figure 4H&I**). The task evoked increase in NE levels in mPFC were different across the four trial outcomes (ANOVA(outcome), F(3)=31.2, p<0.0001), with NE increase in action trials being higher than in withheld trials (p<0.0001, Wilcoxon signed-rank test) (**Figure 4J**). As in S1 reaction time was not dependent on baseline NE levels in mPFC (**Figure 4K**, ANOVA(baseline NE quintile), F(4)=0.12, p=0.97), nor was detection probability (ANOVA(baseline NE quintile x stimulus intensity), F(4,6)=0.95, p=0.46; **Figure 4L**).

Due to the strong relationship between pupil area and behavior, and lack thereof between baseline NE and behavior, we next investigated the relationship between NE dynamics and baseline pupil area (**Figure 5**). NE dynamics differed significantly across baseline pupil area quintiles on hit trials in S1 (ANOVA(baseline pupil quintile x time), F(4,599)=18.9, p<0.0001; and in mPFC (ANOVA(baseline pupil quintile x time), F(4,599)=15.2, p<0.0001; **Figure 5A**). In particular, the lower the baseline pupil area, the higher NE remains elevated following reward delivery, with NE reaching at or below baseline levels by 2 s following the stimulus when pupil area was at its highest. Lick frequency on hit trials did not vary significantly across baseline pupil quintiles (ANOVA(baseline pupil area quintile x time), F(4,59)=1.18, p=0.16; **Extended Data Figure 5-1**), which suggests that elevated NE is related to reward processing rather than motor activity.

**Figure 5.**
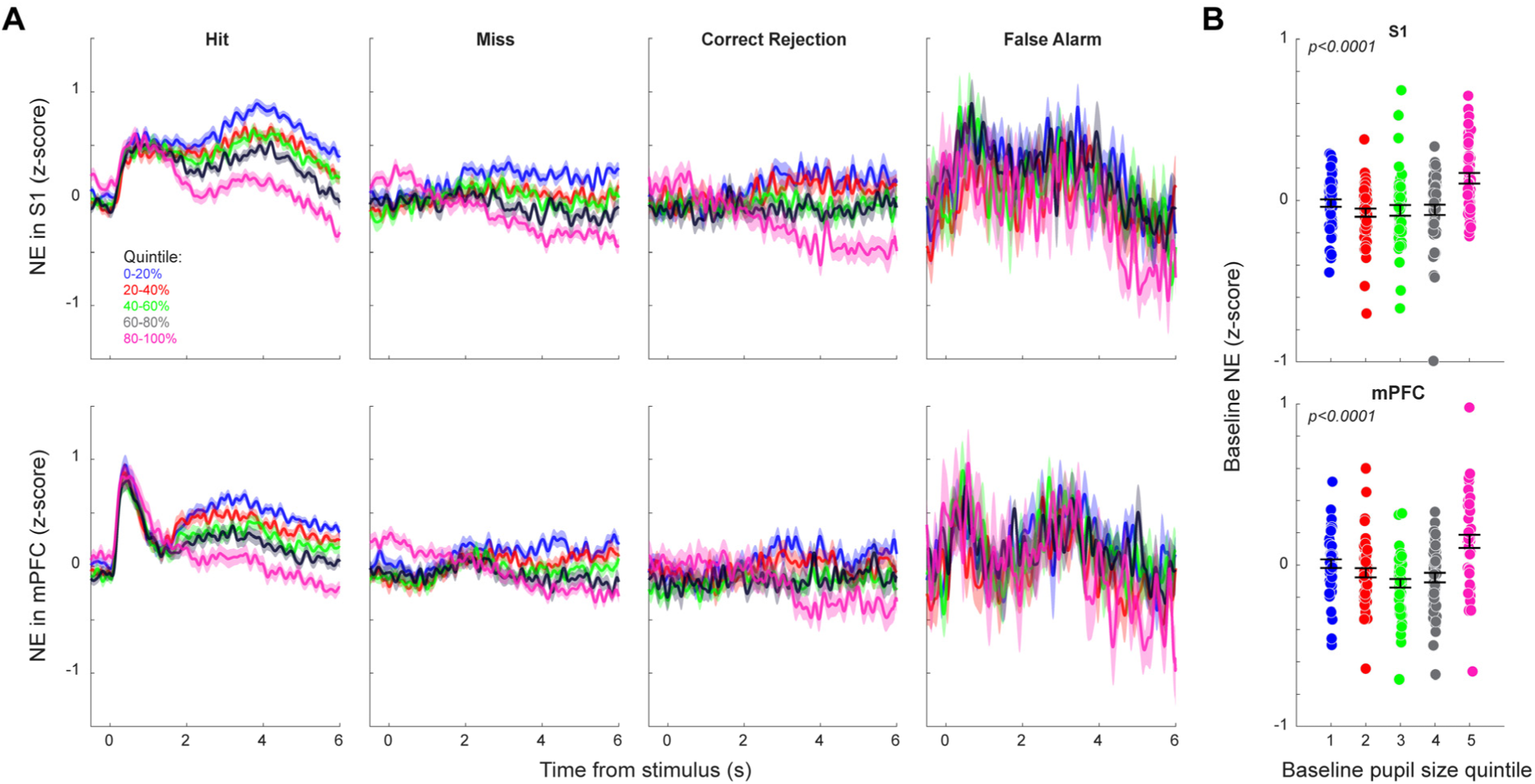
NE dynamics varied strongly with pupil-linked arousal. **A)** Task-evoked NE dynamics in S1 (top) and mPFC (bottom) across trial outcomes and baseline pupil size quintiles. Data here were not separated by stimulus amplitude. Licking dynamics on hit trials did not vary significantly across baseline pupil quintiles (Extended Data Figure 5-1). **B)** Baseline NE levels in S1 (left) and mPFC (right) across trials in each baseline pupil size quintiles. Individual data points represent session average metrics, and error bars represent standard error of mean.

Additionally, baseline NE levels varied significantly with baseline pupil area (**Figure 5B**) in S1 (ANOVA(baseline pupil area quintile), F(4)=9.04, p<0.0001) and in mPFC (ANOVA(baseline pupil area quintile), F(4)=10.9, p<0.0001). Baseline NE levels in both regions followed a left shifted U-shape with the highest baseline NE levels associated with the highest baseline pupil areas, and thus, the lowest task performance (**Figure 2J, K**). This indicated an inverted-U shaped relationship between tonic NE level in S1 and detection performance (Wekselblatt and Niell, 2015).

### GLM-HMM modeling of behavioral state

Recent work has revealed that mice switch between several behavioral strategies or states that are well characterized by hidden Markov models during perceptual decision-making tasks (Ashwood et al., 2022). In our previous analyses, we have shown differences in pupil and NE dynamics depending on trial outcomes (**Figures 2 - 4**), and different NE dynamics depending on pupil-linked arousal on a trial-by-trial basis (**Figure 5**). We therefore hypothesized that distinct behavioral states were associated with distinct NE and pupil dynamics and used GLM-HMM models to uncover the behavioral states that the mice were in during this detection task (**Figure 6**).

**Figure 6.**
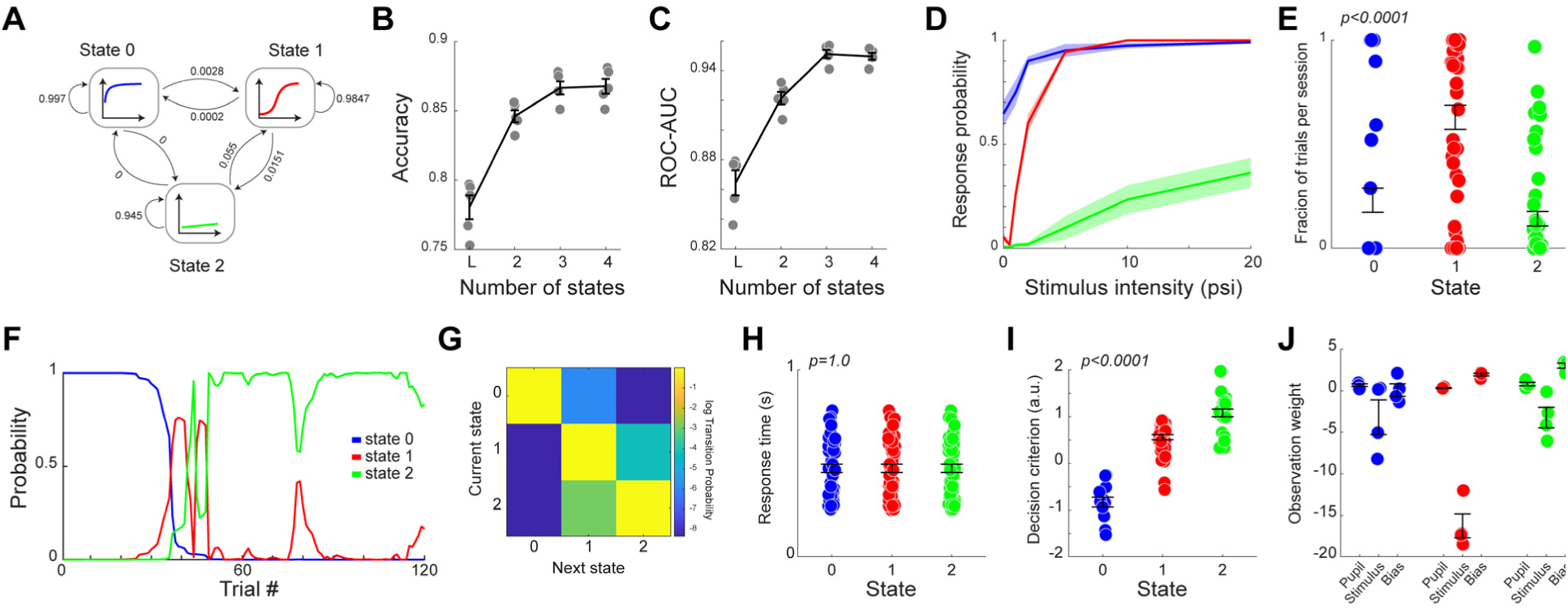
GLM-HMM modeling of behavior. **A)** Diagram of the GLM-HMM model. **B)** Prediction accuracy vs. number of states from 5-fold cross validation. **C)** Area under the receiver operating characteristic curve vs. number of states from 5-fold cross validation. **D)** Average psychometric curve during the three states. **E)** Fraction of total trials during each state per session. **F)** Posterior state probabilities during an example portion of a session. **G)** Inferred transition matrix for the four-state GLM-HMM model. **H)** Reaction time during the four states. **I)** Decision criterion during the three states. **J)** Observation weights for pupil area (used to drive state transitions) and stimulus intensity and bias (used for predicting actions within each state) learned for each animal. Individual data points represent session average metrics unless otherwise noted. Error bars represent standard error of mean.

We employed the GLM-HMM framework from Hulsey et al. (2024) to predict behavioral responses to tactile stimuli where transitions between behavioral states were driven by changes in pupil-linked arousal (see **Methods**). In line with previous work, we compared the prediction accuracy of the lapse model and GLM-HMM models with different numbers of states. A 3-state GLM-HMM had an optimal tradeoff between model accuracy and risk of over-fitting (Ashwood et al., 2022; Le et al., 2023; Hulsey et al., 2024) (**Figure 6B, C**). We also tested whether adding cortical NE signals (baseline mPFC and S1 GRAB_NE_) as additional transition predictors improved the model. Cross-validated performance was statistically indistinguishable from the pupil-only model across K=2–4 (**Extended Data Figure 6-1**; log-likelihood ratio test, p=0.14), so we report the pupil-only transition model throughout. The three states could be characterized as high sensitivity (state 0), intermediate sensitivity (state 1), and low sensitivity or disengaged (state 2) (**Figure 6D**). Mice spent the most trials per session in state 1 and spent the fewest trials in state 2 (**Figure 6E**, ANOVA(state), F(2)=25.2, p<0.0001). Behavioral states were very stable, lasting tens of trials to entire sessions (**Figure 6F, G**). Mice most often alternated between 2 states during a single session (23/49 sessions) Reaction times did not differ across states (ANOVA(state), F(2)<0.0.001, p=1; **Figure 6H**). Decision criterion changed significantly across states; mice had the highest decision criterion (i.e. most conservative) during the low performance state 2 and the lowest decision criterion (i.e. most liberal) during the high performance state 0 (ANOVA(state), F(2)=117.7, p<0.0001) (**Figure 6I**). States were consistent across animals as shown by the weights of models tuned for each mouse (**Figure 6J**).

### Behavioral state dependent pupil and NE dynamics

Since baseline pupil-linked arousal levels drove state transitions in the GLM-HMM, we expected that baseline and task-evoked pupil dynamics would vary depending on behavioral state (**Figure 7**). As expected, baseline pupil area varied significantly between states (ANOVA(state), F(2)=25.3, p<0.0001). Interestingly, baseline pupil area only varied across trial outcomes in the high stimulus sensitivity state 0 (ANOVA(outcome), F(3)=8.4, p<0.001), and not the other two states. Furthermore task-evoked pupil dynamics did not differ significantly across states on hit (ANOVA(state x time), F(2,599)=0.34, p=1) or false alarm trials (ANOVA(state x time), F(2,599)=1.22, p=0.11) but did differ significantly across states on miss (ANOVA(state x time), F(2,599)=2.42, p<0.0001) and correct rejection trials (ANOVA(state x time), F(2,599)=1.32, p=0.043).

**Figure 7.**
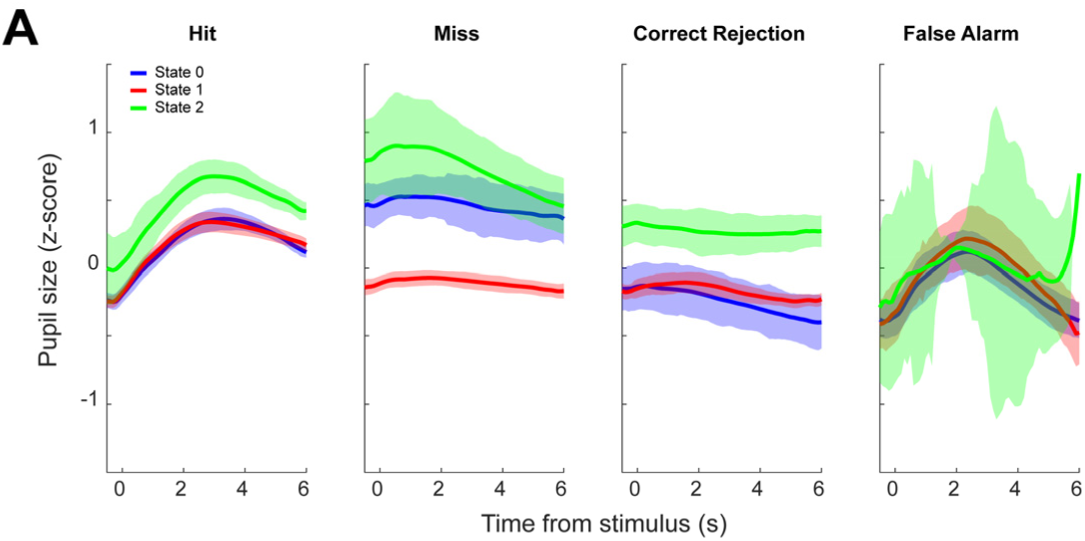
Distinct behavioral state-dependent pupil dynamics. **A)** Task-evoked pupil dynamics in hit, miss, correct rejection and false alarm trials regardless of stimulus amplitude during the three states. Error bars represent standard error of mean.

We also hypothesized that cortical NE dynamics would depend on behavioral state. As expected from our previous analysis (**Figure 3**), baseline synchronization of NE dynamics between S1 and mPFC differed significantly between states (**Figure 8**). Specifically, there was significantly higher shuffle-corrected cross correlation between NE in S1 and mPFC during the disengaged state compared to the other two engaged states (ANOVA(state x lag), F(2,956)=9.18, p<0.0001). While there were no significant differences in baseline NE in either mPFC or S1 across states, we observed distinct task-evoked NE dynamics in S1 and mPFC. For instance, NE dynamics differed significantly across states in S1 (ANOVA(state x time), F(2,599)=1.30, p<0.0001) and mPFC (ANOVA(state x time), F(2,599)=1.82, p<0.0001).

**Figure 8.**
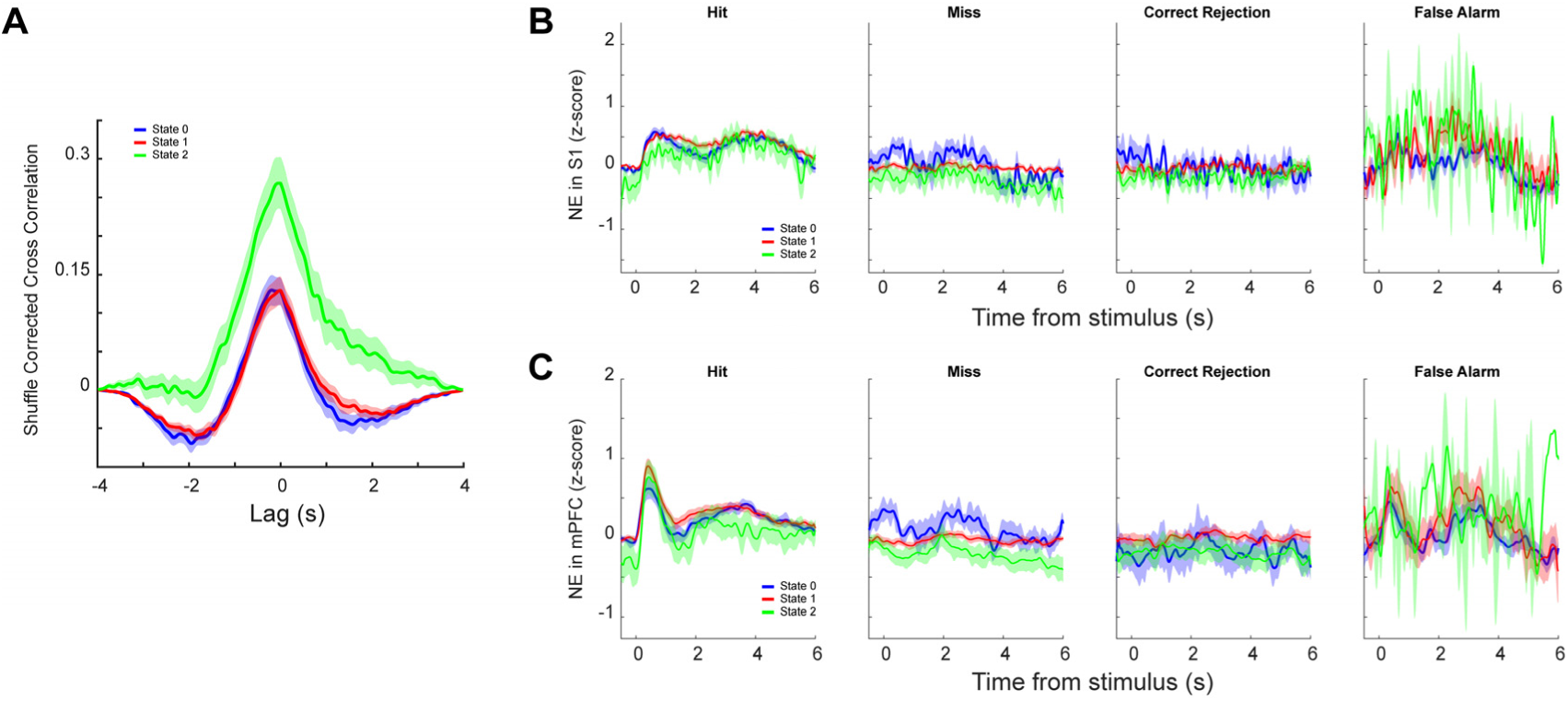
Distinct behavioral state-dependent NE dynamics in S1 and mPFC. **A)** Average cross-correlation between baseline NE signals in S1 and mPFC regardless of stimulus amplitude during the three states. **B)** Task-evoked NE dynamics in S1 in hit, miss, correct rejection and false alarm trials during the three states. **C)** Task-evoked NE dynamics in mPFC in hit, miss, correct rejection and false alarm trials during the three states. Error bars represent standard error of mean.

## Discussion

In this study we trained mice to perform a whisker-based tactile signal detection task while simultaneously monitoring pupil dynamics and NE activity in S1 and mPFC using GRAB_NE_ fiber photometry. We found that both pupil size and synchronization of cortical NE signals were tightly linked to whether the animal reported the stimulus (hit and false alarm trials) versus withheld a response (miss and correct rejection trials), but that baseline pupil size and NE dynamics carried distinct information about behavioral states and trial outcomes. Together, these results support a nuanced view in which pupil size and cortical NE are related but not interchangeable readouts of arousal and engagement (Joshi and Gold, 2020; Cazettes et al., 2021; Liu et al., 2021; Yang et al., 2021; Megemont et al., 2022; Grujic et al., 2024; Weiss et al., 2025).

Consistent with previous work in rodents, we observed large, reliable pupil dilations following stimulus onset and during licking on hit trials, as well as similarly robust dilations on false alarm trials, with minimal changes on miss and correct rejection trials (**Figure 2**). The latency from stimulus onset to pupil dilation in our detection task was relatively short (on the order of ∼100 ms), shorter than reported in whisker-based discrimination paradigms that require texture or direction judgments, where dilation emerges later (Lee and Margolis, 2016; Schriver et al., 2018). In addition, baseline pupil size also differed in our mice compared with a previous study in rats. Mice showed larger baseline pupils on trials in which they subsequently withheld a response (miss and correct rejection) and smaller baseline pupils on trials in which they responded (hit and false alarm). Although this finding supports the use of baseline pupil size as an index of arousal or engagement, it runs opposite to what we previously observed in rats performing a more complex tactile discrimination task, where baseline pupil size tended to be higher on response trials (Schriver et al., 2018). While this may reflect some genuine difference between species, these differences are likely related to task demands: detection requires only deciding whether a stimulus is present, whereas discrimination requires additional sensory evaluation and decision processes that may delay the arousal-related pupil response.

Using GRAB_NE_, we observed rapid, stimulus-locked increases in NE in both cortical regions on hit and false alarm trials, and negligible changes on miss and correct rejection trials, mirroring the trial outcome dependence of the pupil response (**Figure 4**). However, the temporal profiles of NE signals were distinct between S1 and mPFC. On hit trials, NE in mPFC showed a sharp transient peak followed by a broader, lower-amplitude elevation, whereas NE in S1 exhibited two broader peaks of comparable magnitude. These distinct temporal patterns suggest that noradrenergic signaling from the LC were not broadcast uniformly to all cortical targets; rather, distinct subpopulations of noradrenergic neurons in the LC project with distinct task-related spiking dynamics project to different cortical areas (Totah et al., 2018; Breton-Provencher et al., 2021; Kelberman et al., 2024). In support of this notion, recent work has shown that distinct subpopulations in LC project to basal forebrain and mPFC with only 30% overlap in neurons that project to both (Liu et al., 2025).

The feedback from mPFC to LC may shape this divergence in NE dynamics (Poe et al., 2020; Totah et al., 2021; Kelberman et al., 2024). mPFC provides the densest cortical input to LC and has been proposed to use LC as a hub to broadcast control signals to other brain regions (Jodoj et al., 1998; Mashour et al., 2022). Stimulation of mPFC can elicit LC spiking, whereas stimulation of more lateral frontal regions is less effective, suggesting a privileged mPFC–LC pathway (Jodoj et al., 1998). In this framework, excitatory projections from mPFC onto inhibitory LC interneurons could transiently suppress LC output back to mPFC, producing a sharp peak followed by a rapid decline in NE in mPFC (Breton-Provencher and Sur, 2019). At the same time, excitatory projections from mPFC onto LC noradrenergic neurons that project to S1 could maintain a more sustained elevation of NE in S1, consistent with the longer-lasting S1 responses that we observed (**Figure 4**). Alternatively, noradrenergic projections from LC to mPFC may exhibit increased expression of presynaptic inhibitory α2 receptors compared to those that project to S1. Although our data are correlational, this model provides a testable circuit-level explanation for the observed differences between mPFC and S1.

In contrast to baseline pupil size, baseline NE levels in S1 and mPFC alone did not predict whether the animal would respond on the trial. Instead, we observed asymmetric U-shaped relationships between baseline pupil area and baseline cortical NE levels (**Figure 5B**). We also found that the correlation of GRAB_NE_ signals between S1 and mPFC during the baseline period was higher on withheld trials than on response trials (**Figure 3**). If LC contains partially segregated subpopulations projecting to S1 and mPFC, greater baseline synchrony between these projections may reflect a more homogeneous, low-information state of noradrenergic drive during behavioral disengagement (Wang et al., 2010). In contrast, lower cross-correlation before response trials could indicate more differentiated, target-specific NE signaling when the animal is engaged and prepared to report stimuli. Thus, whereas baseline pupil area appears to index a global arousal or engagement, the baseline coordination of NE across cortical targets may capture a more circuit-specific facet of the animal’s internal state.

We also found that pupil-linked arousal strongly modulated cortical NE dynamics (**Figure 5A, B**). High baseline pupil area and low task performance were associated reduced cortical NE following reward on hit trials. Cortical NE levels following reward were highest when baseline pupil area was lowest. As we did not observe any accompanying relationship between motor activity following reward and baseline pupil area (**Extended Data Figure 5-1**), our findings suggest that processing of rewards in LC, and consequently in the cortical targets of NE release, is strongly modulated by arousal.

Finally, we used the GLM-HMM framework developed by Ashwood et al. (2022) and expanded on by Hulsey et al. (2024) with transitions between behavioral states driven by pupil-linked arousal to model behavior on a Go/No-Go task (**Figure 6**). As in previous work (Ashwood et al., 2022; Le et al., 2023; Hulsey et al., 2024), task performance was best described with three states: here corresponding to a high sensitivity state with high stimulus detection and a high false alarm rate, an intermediate state with near optimal performance, and a low sensitivity or disengaged state. We observed distinct task-related pupil and NE dynamics across the three states (**Figure 7, 8**). These differences in dynamics suggest that the latent states identified in the GLM-HMM framework corresponded to genuine brain states with distinct computations underpinning stimulus detection, decision making, and reward processing. Future work is warranted to elucidate how NE differentially modulates task-relevant neural information processing across these latent states.

This study did have several limitations, including a small number of exclusively male mice (4), a narrow scope of cortical regions interrogated through fiber photometry (S1 and mPFC), and a relatively simple task (signal detection). Further studies may investigate whether the relationships between pupil dynamics, cortical NE release, and behavioral state shown here hold in more cognitively demanding tasks like signal discrimination and across sensory modalities (e.g. auditory or visual detection/discrimination). Additionally, a more complete understanding of noradrenergic dynamics across cortical regions may be obtained using wide-field imaging.

## Acknowledgements

This work was supported by NIH R01NS119813, R01AG075114, R21MH125107, and NSF CBET 1847315.

## Data availability

All data is available upon request. All code is available at https://github.com/Neural-Control-Engineering/Neurodynamic-control-toolbox

## Disclaimer

Q.W. is a co-founder of Sharper Sense.

## Extended Data

**Extended Data Figure 2-1.**
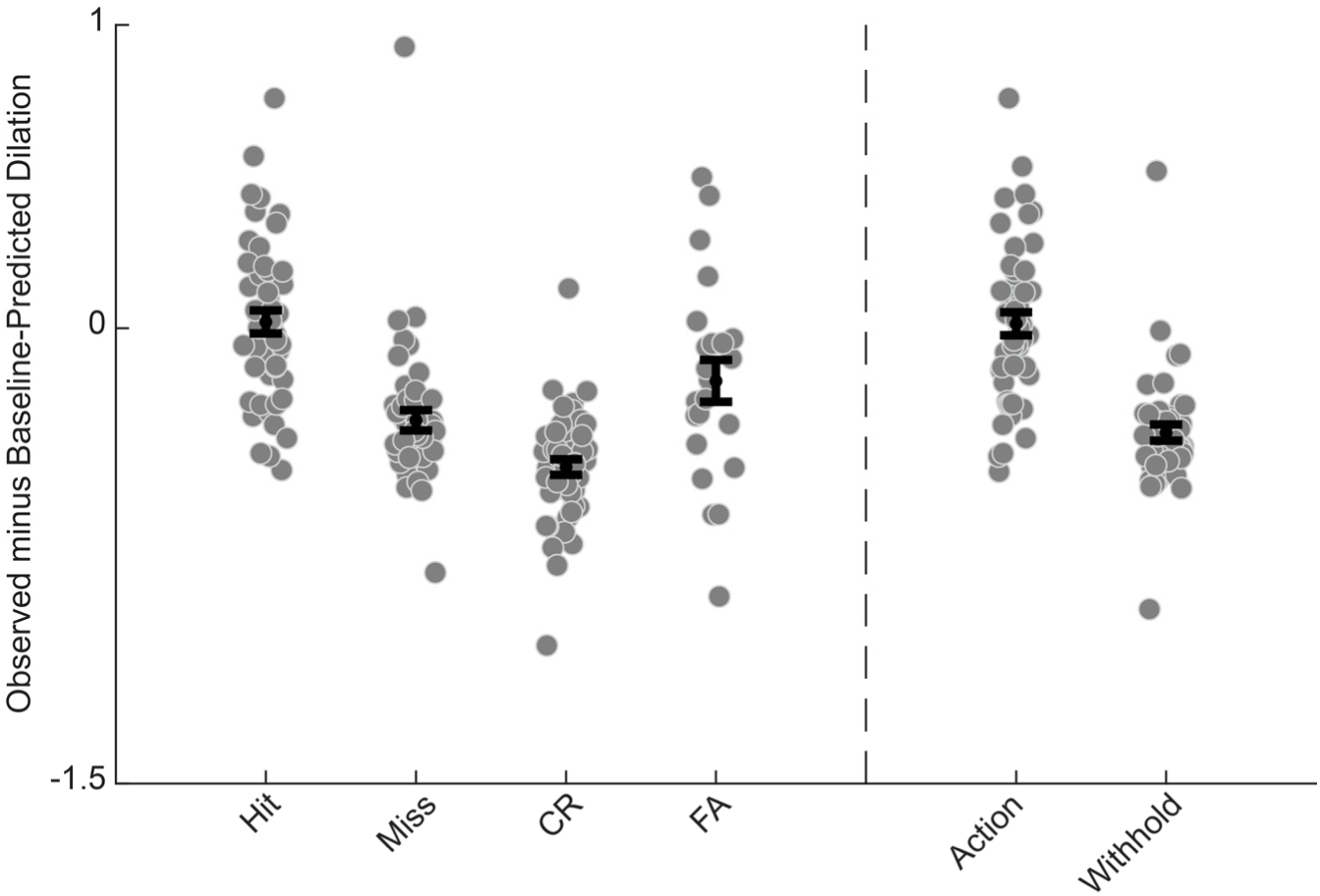
Baseline-corrected pupil dilations across trial outcomes. Individual data points are session average baseline-corrected dilations for a given outcome. Error bars represent standard error.

**Extended Data Figure 5-1.**
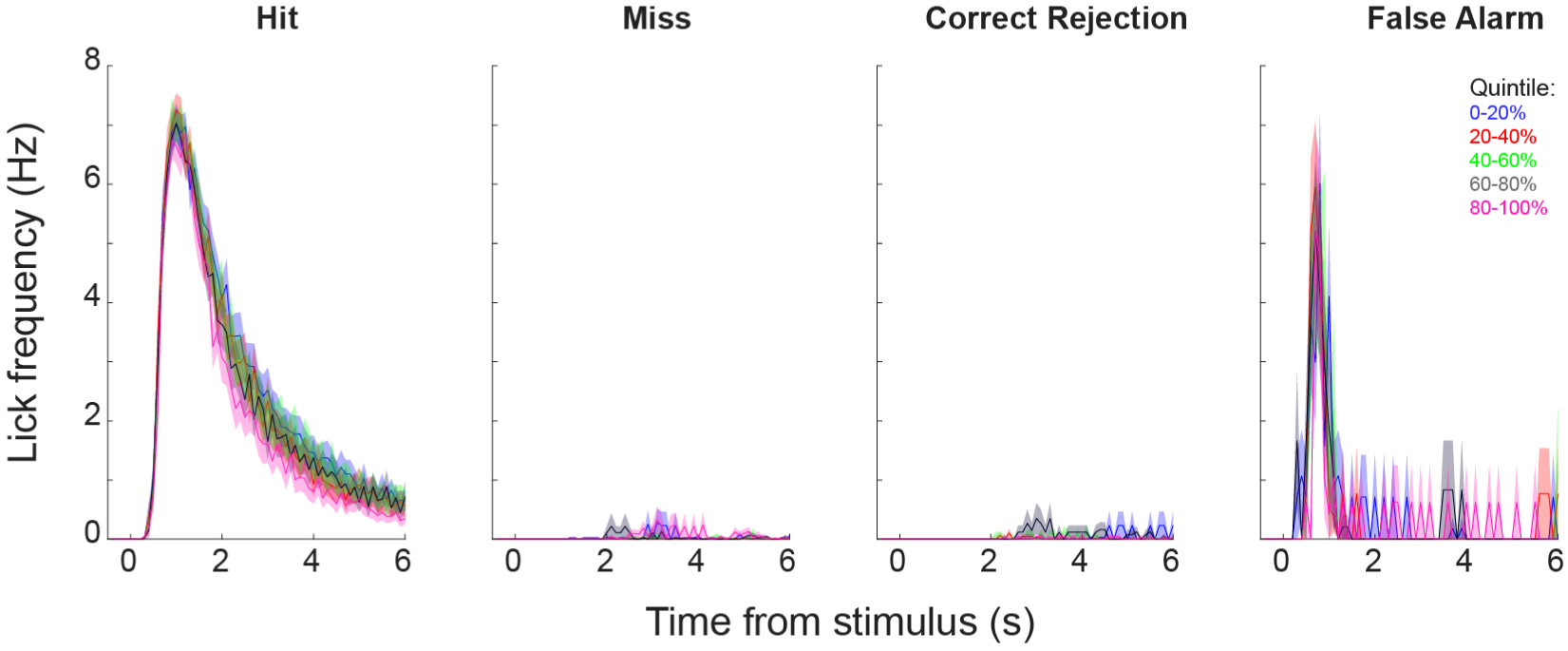
Task-evoked lick frequency separated by baseline pupil area quintile and task outcome. Error bars represent standard error.

**Extended Data Figure 6-1.**
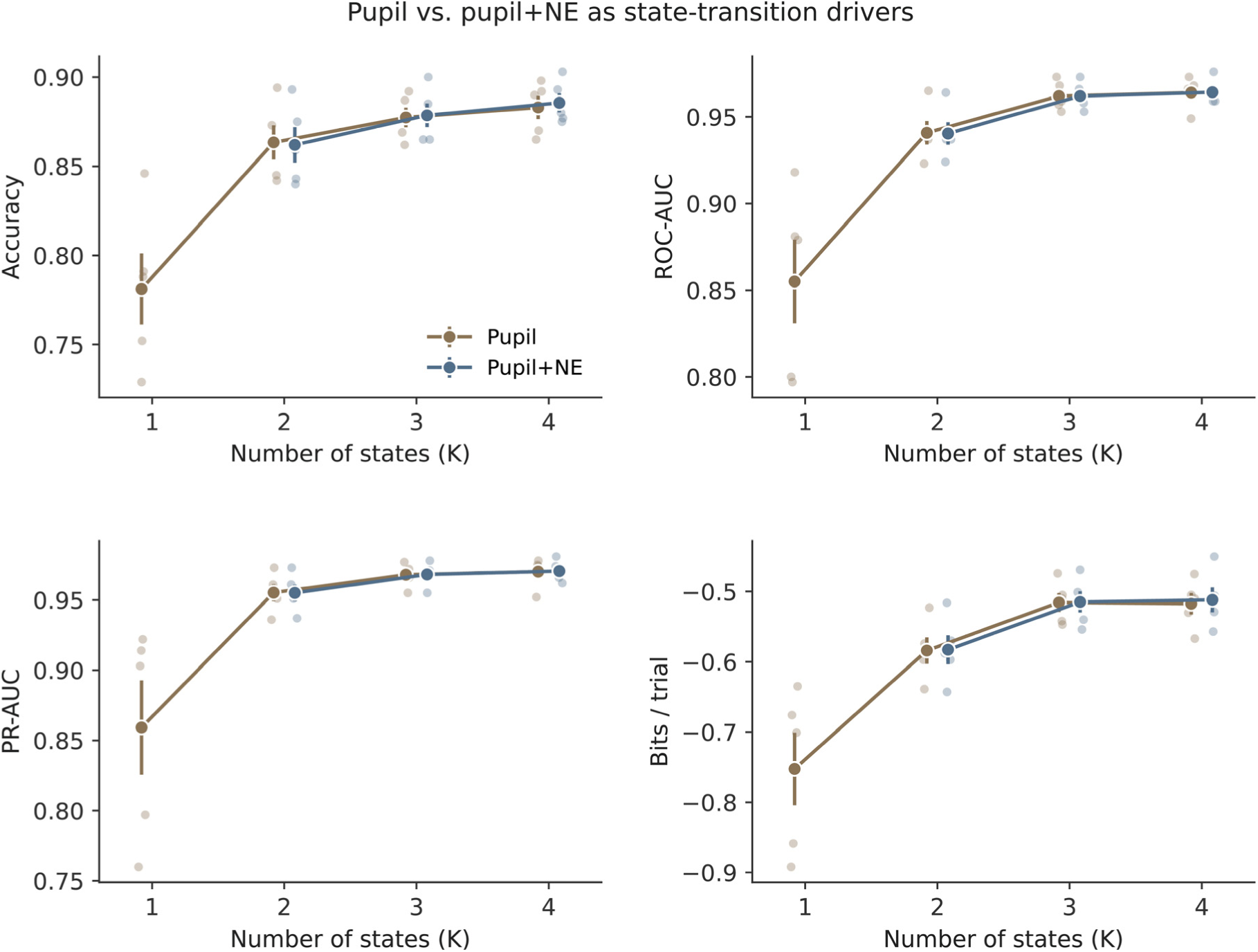
Comparison of A) predictive accuracy, B) area under the receiver operant characteristic curve, C) area under the precision-recall curve, and D) bits/trial across states between a GLM-HMM where state transitions were driven by baseline pupil area alone and a GLM-HMM where state transitions were driven by baseline pupil area and baseline NE in mPFC and S1. Individual data points are from 5-fold cross validation, error bars represent standard error of mean.

